# Structurally conserved primate lncRNAs are transiently expressed during human cortical differentiation and influence cell type specific genes

**DOI:** 10.1101/232553

**Authors:** Andrew R. Field, Frank M.J. Jacobs, Ian T. Fiddes, Alex P.R. Phillips, Andrea M. Reyes-Ortiz, Erin LaMontagne, Lila Whitehead, Vincent Meng, Jimi L. Rosenkrantz, Maximillian Haeussler, Sol Katzman, Sofie R. Salama, David Haussler

## Abstract

The cerebral cortex has expanded in size and complexity in primates, yet the underlying molecular mechanisms are obscure. We generated cortical organoids from human, chimpanzee, orangutan, and rhesus pluripotent stem cells and sequenced their transcriptomes at weekly time points for comparative analysis. We used transcript structure and expression conservation to discover thousands of expressed long non-coding RNAs (lncRNAs). Of 2,975 human, multi-exonic lncRNAs, 2,143 were structurally conserved to chimpanzee, 1,731 to orangutan, and 1,290 to rhesus. 386 human lncRNAs were transiently expressed (TrEx) and a similar expression pattern was often observed in great apes (46%) and rhesus (31%). Many TrEx lncRNAs were associated with neuroepithelium, radial glia, or Cajal-Retzius cells by single cell RNA-sequencing. 3/8 tested by ectopic expression showed ≥2-fold effects on neural genes. This rich resource of primate expression data in early cortical development provides a framework for identifying new, potentially functional lncRNAs.

## INTRODUCTION

Advances in pluripotent stem cell technology have allowed researchers to use organoid technology to probe gene regulatory events that occur during the differentiation of early neocortical cell types using cell lines that model both normal and disease states (Eiraku et al., 2008; Eiraku and Sasai 2012; Lancaster et al., 2013; Qian et al 2016). These protocols have been shown to closely recapitulate cellular organization and gene expression events observed in fetal tissue (Camp et al., 2015; reviewed in Fatehullah et al., 2016). Comparisons of human with primate organoids have revealed subtle differences in the timing of cell divisions and differentiation events (Mora-Bermudez et al., 2016; Otani et al., 2016), though the mechanisms by which these changes are enacted are unknown.

Here we focus on one class of gene regulatory element, long non-coding RNAs (lncRNAs), which often show tissue specific expression (Ponting et al., 2009; Cabili et al., 2011; Derrien et al 2012; Pauli et al., 2012), account for a significant proportion of Pol II output (Carninci et al 2005; Harrow et al., 2006; Derrien et al., 2012), and show particular enrichment in neural tissues (Ravasi et al., 2006; Cabili et al., 2011; Derrien et al., 2012; Ramos et al., 2013). LncRNAs are demonstrated to have diverse roles in gene regulation including chromosome inactivation (Penny et al., 1996; Zhao et al., 2008), imprinting (Lighton et al., 1995; Camprubi et al., 2006; Buiting et al., 2007; Pandey et al., 2008; Martins-Taylor et al., 2014), and developmental processes (Rinn et al., 2007; Heo and Sung, 2011) and have been implicated in establishment of pluripotency (Guttman et al., 2009; Guttman et al., 2011), stem cell maintenance (Rani et al., 2016), reprogramming (Loewer et al., 2010), and differentiation (Guttman et al., 2011). Nevertheless, most of the tens of thousands of identified lncRNAs in human have undetermined function (Hon et al., 2017; Lagarde et al., 2017) and lack sequence conservation among vertebrate species (Wang et al., 2004; Church et al., 2009; Cabili et al., 2011; Ulitsky et al., 2011; Kutter et al., 2012). Their tissue specific expression patterns, rapid sequence evolution, and implication in cell type specification make lncRNAs an attractive target as arbiters of species-specific gene regulation during development.

It has been suggested that exon structure conservation is a better indicator of conserved function than nucleic acid sequence alone (Ulitsky, 2016) and we postulate that expression pattern conservation during differentiation may imply a conserved role in cell type specification. Here, we utilized an approach focusing on both of these aspects of conservation in equivalent developing tissues among closely related primate species to identify gene regulatory lncRNAs active in human neural differentiation. We generated cortical organoids from human, chimpanzee, orangutan, and rhesus pluripotent stem cells to recapitulate early events in cortical development and enable comparative molecular analysis of this process. RNA-sequencing was performed on weekly time points to assess the conservation of lncRNA transcript structure and expression among primates. Particular attention was paid to lncRNA transcripts that were transiently expressed (TrEx lncRNAs), specifically those with max expression after neural induction but diminished expression by week 5 of our protocol, for their implication in early neurogenesis and their potential to contribute to human or primate-specific attributes of the cerebral cortex. Single cell RNA-sequencing on a subset of time points relevant to major differentiation events was used to identify the cell subpopulations associated with the expression of candidate TrEx lncRNAs. Finally, CRISPR activation (CRISPRa) in HEK293FT cells was used to express these transcripts out of context to probe whether transiently expressed lncRNAs can regulate the gene networks associated with them in our single cell data. In all, we identified 8 TrEx lncRNAs whose expression had repeatable effects on protein-coding genes correlated or anticorrelated with the transcripts in single cell data. Though most were modest effects, 3 lncRNAs, TrEx108, TrEx5008, and TrEx6168, showed a significant induction or reduction of correlated genes by 2-fold or more.

## RESULTS

### Generation and RNA-sequencing of primate cortical organoids

To study the transcriptional landscape of early cell type transitions during primate cortical neuron differentiation, we subjected human, chimpanzee, orangutan, and rhesus macaque pluripotent stem cells to a cortical neurosphere differentiation protocol based on Eiraku et al., 2008 and optimized for use with our cell lines (Fig. 1A & B). Embryonic stem cell lines were used for human (H9) and rhesus (LYON-ES1) time courses, but, since embryonic stem cells are not available for great apes, we generated integration-free induced pluripotent stem cell lines for chimpanzee (Epi-8919-1A) and orangutan (Jos-3C1) from primary fibroblasts (Fig. S1 & S2).

**Figure 1.**
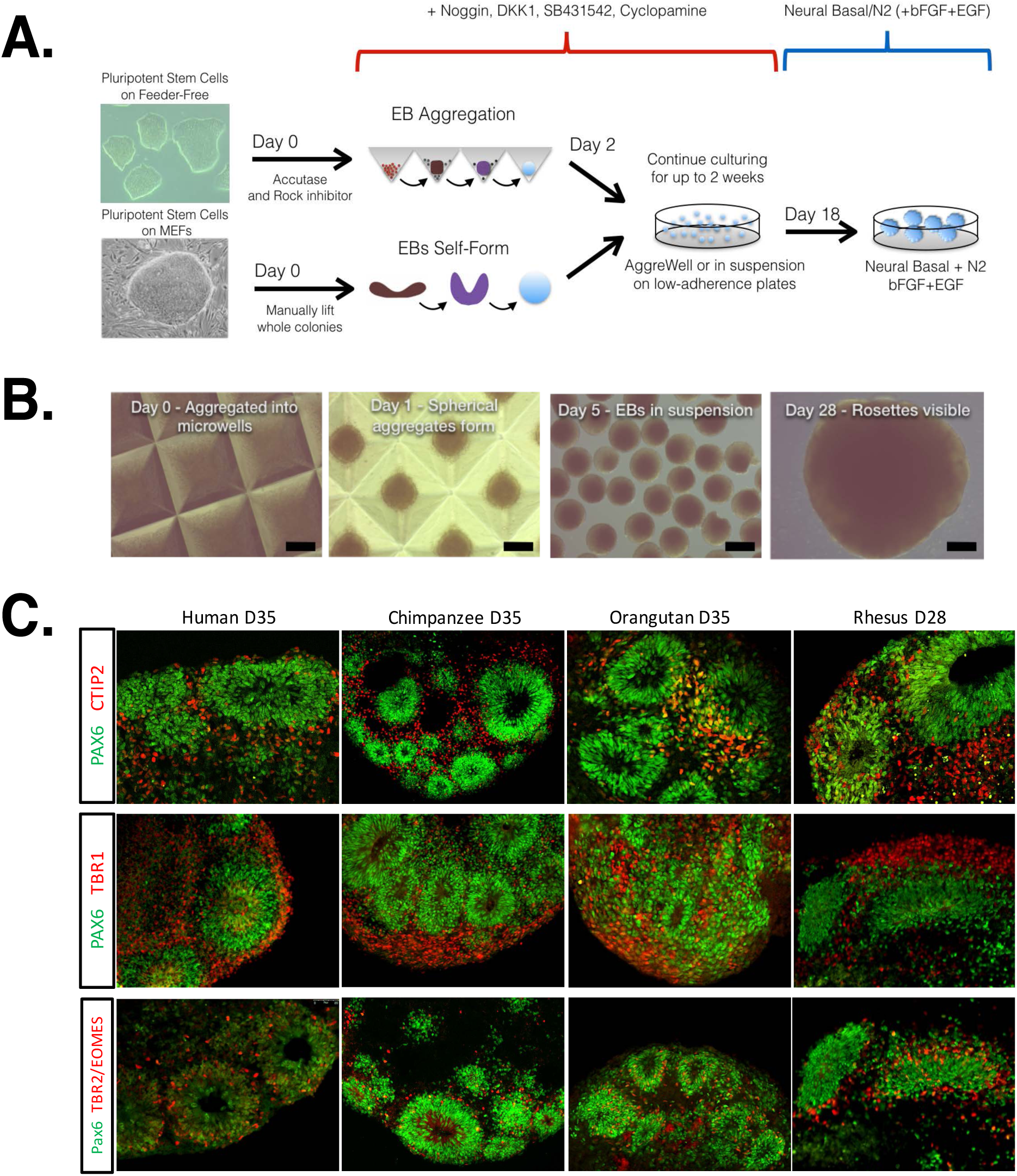
Cortical neural epithelium differentiation protocol. (A) An outline of our dorsal neuron differentiation assay for both feeder grown and feeder-free cultures is provided. Cells grown on MEF feeders were manually lifted and allowed to self-form into embryoid bodies. Cells in feeder-free conditions were first dissociated into single cell suspensions and aggregated into embryoid bodies consisting of approximately 10,000 cells using AggreWell plates (Stem Cell Tech). Embryoid bodies were then incubated with the addition of Noggin, DKK1, SB431542, and cyclopamine for 18 days to induce cortical neuron differentiation. The cultures were then switched to Neural Basal media supplemented with N2 with the addition of bFGF and EGF for feeder-free cultures. (B) An example of chimpanzee aggregation and differentiation at days 0, 1, 5, and 28 is shown. (C) Immunofluorescence staining at our endpoint of 5 weeks (28 days in rhesus). PAX6, marking neural progenitors, form the center of a rosette pattern forming around lumen like structures that are prominent throughout neurospheres in each species. CTIP2 and TBR1 indicate early deep layer neurons projected radially out from neural progenitors. Finally, TBR2 is characteristic of intermediate progenitors or early migrating neurons as they transit radially outward from the lumen-like structures.

Performance of these stem cell lines in our cortical organoid differentiation assay was evaluated by immunofluorescence staining at day 35 (or the equivalent day 28 in rhesus; see Methods) showing efficient production of radial glia, intermediate progenitors, and early deep layer cortical neurons (Fig. 1C) in highly structured neural rosettes as described previously (Eiraku et al., 2008; Eiraku and Sasai, 2012; Lancaster et al., 2013; Camp et al., 2015; Qian et al., 2016). RNA samples were collected in at least two replicates from pluripotent stem cells and weekly time points over 5 weeks of neural differentiation in each species and used to create total-transcriptome strand-specific RNA sequencing libraries. Due to their shorter gestational period and the faster division rates of their embryonic stem cells, rhesus macaque samples had adjusted time points with ~5.5 day weeks (Fig. S3; Methods). In all, over 2 billion paired-end RNA-seq reads were uniquely mapped to their respective genomes from 49 libraries, averaging 41 million reads per library with a minimum of 46 million total reads across replicates per species time point. After mapping to the appropriate genome and gene annotation file (Fig. 2A; Methods) DESeq (Love et al., 2014) was used to assess relative gene expression for known genes. The differentiation protocol is designed to generate primarily dorsal cortical tissue. Dorsalization was confirmed by the transcriptional profile of dorsal marker genes in all species (Fig. 2B). Pluripotency markers such as OCT3/4 are downregulated by the week 1 time point in all species and early neural stem cell markers such as PAX6 are upregulated. There was strong expression of deep layer neuron markers, such as TBR1, by week 5 in all species (Fig. 2C). Overall, there is strong induction of early neural differentiation and dorsal forebrain markers with little expression of markers of other brain regions (Fig. 2B).

**Figure 2:**
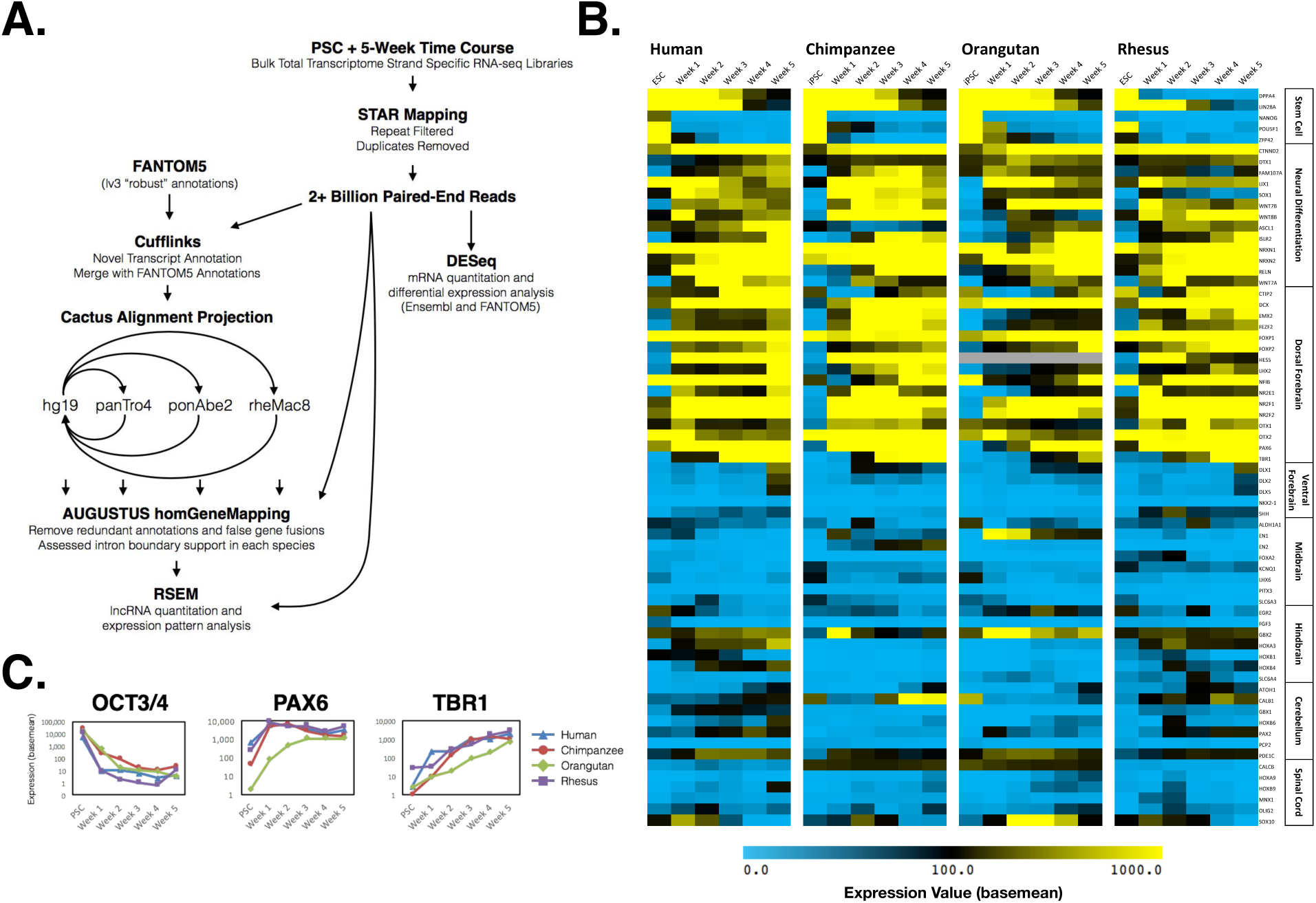
Transcriptomic analysis and marker gene expression. (A) A flowchart depicts the analysis pipeline used to process the neural differentiation time course RNA-seq data and identify expressed lncRNAs. (B) A heatmap with gene expression colored by basemean values calculated from DESeq2. Canonical marker genes for pluripotent stem cells, neural differentiation, dorsal forebrain, ventral forebrain, midbrain, hindbrain, cerebellum, and spinal cord are displayed. (C) Plots depicting the expression in basemean value of the genes *OCT3/4, PAX6,* and *TBR1* in human (blue), chimpanzee (red), orangutan (green), and rhesus (purple) over the neural differentiation time course.

To assemble potential novel transcripts in each species, Cufflinks v2.0.2 (Trapnell et al., 2010; Trapnell et al., 2012) was used and the Cuffmerge tool combined gene models across time points in each respective species using FANTOM5 lv3 (Hon et al., 2017) as a reference annotation. CAT (Fiddes et al., submitted) was used to project the FANTOM lv3 set through a progressive Cactus whole-genome alignment (Stanke et al., 2008; Paten et al., 2011) to each of the other primate genomes (Fig. 2A). Guided by the Cufflinks annotation set in each genome, these projections from the other genomes were assigned a putative gene locus. RSEM v1.3.0 (Li and Dewey, 2011) was used to calculate expression values of these new gene models (Fig. 2A).

### Expression and gene structure conservation of lncRNAs across primates

It has been suggested that conservation of exon boundaries within a lncRNA gene may be a feature indicative of functional transcripts (Ulitsky, 2016). Gene structure conservation of expressed transcripts among our primate species was assessed using the homGeneMapping tool from the AUGUSTUS toolkit (Konig et al., 2016). homGeneMapping makes use of cactus alignments to project annotation features in all pairwise species comparisons, providing an accounting of features found in other genomes. homGeneMapping was provided with both the Cufflinks transcript assemblies as well as expression estimates derived from the combination of the week 0 to week 5 RNA-seq experiments in all four species. The results of this pipeline were combined with the Cactus alignment-based transcript projections to ascertain a set of gene loci that appear to have human-specific expression, human-chimp specific expression, great ape-specific expression, and expression in all primates (Fig. 3A & B, Table S1). Transcript models that had at least 50% intron junction support in human were considered conserved in a non-human primate genome if that genome had RNA-seq read support for any of its intron junctions and the gene cluster had a TPM value greater than 0.1. All single-exon transcripts were filtered out to reduce noise.

**Figure 3:**
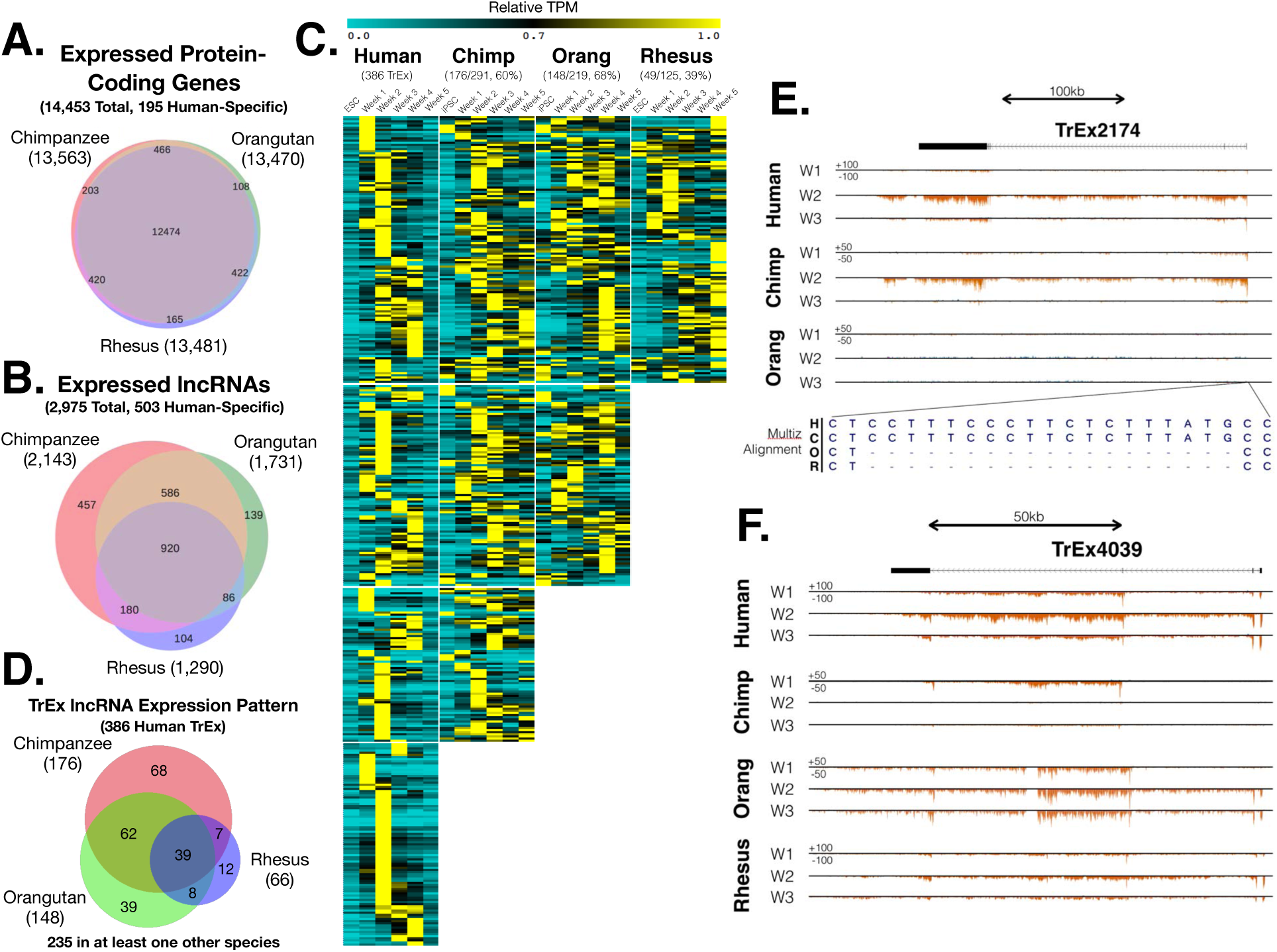
LncRNA structure and expression pattern conservation. Venn diagrams show the degree of intron boundary conservation of human protein-coding genes (A) and lncRNAs (B) in each of the other species. LncRNAs that had transient expression patterns in human samples were compared across species (C and D). (C) A heatmap depicting expression in relative TPM normalized to the max value in each species shows expression patterns of human TrEx lncRNAs with conserved structure in the other species. (D) Venn diagram showing TrEx lncRNA expression pattern conservation between species. UCSC Genome Browser screenshots show the expression of TrEx2174 (E) and TrEx4039 (F) in their respective conserved species. The Multiz alignment just upstream of the TSS for TrEx2174 has a 19bp insertion that is specific to human and chimpanzee among extant great apes (E).

Using these parameters, 2,975 human poly-exonic lncRNA gene clusters were identified in our human RNA-seq reads. 503 human-specific, 457 human-chimp specific, 586 great ape-specific, and 920 primate-conserved lncRNAs were found by intron boundary support and a minimum expression of at least 0.1 TPM (Fig. 3B, Table S1). These show higher overlap in species that are separated by less evolutionary distance as expected. Among the primate conserved category are the previously described mammalian conserved lncRNAs *MALAT1*, *NEAT1*, *H19*, *PRWN1*, and *CRNDE* (Table S1). In addition to known lncRNAs, 347 novel gene clusters were also found by Cufflinks in the human RNA-seq data, 160 of which were human-specific, 164 conserved in chimp, 105 in great apes, and 79 conserved across all of the primates (Fig. S4, Table S2), displaying a similar distribution to lncRNAs defined by the FANTOM5 consortium (Hon et al., 2017). 580 chimpanzee-specific, 1,709 orangutan-specific, and 593 rhesus-specific gene loci were also detected (Table S2) further supporting a relatively fast turn-over of transcribed intergenic loci, though we suspect that the orangutan estimates are inflated due to its relatively poor genome assembly and, consequently, poor alignment to the other genomes. Comparing these figures to protein-coding genes, 14,453 coding genes were found to be expressed in human (Table S1), and 12,474 (86%) of these coding genes were expressed and shared intron boundaries among all species (Fig. 3A). This confirms a much higher degree of gene structural conservation of mRNAs by these strict metrics.

LncRNAs have previously been reported to exhibit dynamic expression in developing tissues (Cao et al., 2006; Amaral and Mattick, 2008; Chodroff et al., 2010; Qureshi et al., 2010; Pauli et al., 2012). These transiently expressed (TrEx) lncRNAs offer a promising mechanism for the rapid evolution of regulatory networks in developing tissues, though few, as of yet, have been shown to have gene regulatory function. We looked for lncRNAs expressed transiently in our data set, defining TrEx lncRNAs as those with maximal expression between weeks 1 and 4, and less than 50% their maximal expression at weeks 0 and 5. Using these metrics, we identified 386 human TrEx lncRNAs, most of which were expressed primarily at one weekly time point (Fig. 3C & D, Table S3). We next assessed if these transcripts were also transiently expressed in other species, requiring that they also have max expression at weeks 1-4 in that species. In all, 39 (31% of 125 transcripts with conserved structure in all 4 species) had a transient expression pattern in all 4 species and 235 (81% of 291 transcripts with conserved structure in at least chimpanzee) were TrEx in human and at least one other species. 176 were conserved in chimpanzee (61% of 291 transcripts with conserved structure), 148 in orangutan (68% of 219), and 66 (53% of 125) in rhesus (Fig. 3C & D, Table S3).

Several examples highlight the general features of these TrEx lncRNAs and illustrate their potential evolutionary impact. TrEx2174 (RP11-314P15) is notable in its week 2 specific expression which is also observed in chimpanzee (Fig. 3E), while not being expressed in orangutan or rhesus at any time point. Interestingly, TrEx2174 has a 19 bp insertion which overlaps its transcription start site that is specific to human and chimpanzee. This suggests TrEx2174 may be a very recently evolved lncRNA, or a lncRNA with a recently evolved expression pattern. Among the lncRNAs that were observed in all 4 of our species was TrEx4039 (overlapping AC011306 and MIR217HG) (Fig. 3F). TrEx4039 expression peaks at weeks 1 or 2 in all species and is extinguished by week 5. Chimpanzee appears to express an isoform of this transcript that is not shared with human or rhesus, but can be seen expressed early in orangutan. While chimpanzee ceases expression from this locus at week 2, orangutan appears to switch to the longer isoform observed in the other two species at week 2. This demonstrates how even among transcripts that share structural elements across species, expression regulation can be diverse.

### Transiently expressed lncRNAs show cell type-specific expression patterns

Since bulk RNA-sequencing represents an average gene expression value across multiple cell types present in our organoids, which obscures cell type specific signals, we performed 10X Chromium 3’ end single cell RNA-sequencing on human ESCs and cortical organoids at weeks 0, 1, 2, and 5 to assess the cell type specificity of TrEx lncRNAs. Reads were mapped with the CellRanger 2.0 pipeline using the modified FANTOM5 gene models as for the bulk RNA-Seq. In all, nearly 800 million reads were obtained from 14,086 cells averaging 56.6k reads per cell. The total number of genes detected per library ranged from 28k in week 5 neurospheres to 36k at week 1, averaging between 1702 and 4978 genes per cell.

tSNE plots generated with CellRanger v1.2 (10X Genomics) identified increasing cell heterogeneity as differentiation progressed (Fig. S4). Using a combination of k-means clustering, graphical clustering, and visual inspection, we were able to manually curate clusters of cells with expression profiles matching neuroepithelium (NE), radial glia (RG), and Cajal-Retzius (CR) cells in our week 2 libraries (Fig. S5, Table S4). NE cells were identified by expression of *HES3* and *NR2F1* forming a cluster of 1261 cells (29%) (Fig. S5C). CR cells expressed *TBR1, EOMES, LHX9,* and *NHLH1* comprising a cluster of 356 cells (8%) (Fig. S5E). The largest proportion of cells showed strong expression of cortical RG markers *SOX2, EMX2, NNAT, PTN,* and *TLE4* making up 2593 cells (59%) (Fig. S5D). About 176 cells (4%) showed no strong association with these manually curated cell clusters and had no significant distinguishing genes. We determined that they likely represented cell doublets; their prevalence is consistent with theoretical estimates of the number of doublets from the number of cells we captured per library. At week 5, cells expressing NE markers were virtually absent and instead additional clusters expressing mature neuronal markers emerged (Fig. S4).

LncRNAs have previously been associated with specific tissues and developmental time points (Cabili et al., 2011; Ponting et al., 2009; Derrien et al., 2012; Pauli et al., 2012), including spatial and temporal expression patterns within the brain (Cao et al., 2006; Amaral and Mattick, 2008; Chodroff et al., 2010; Qureshi et al., 2010), and to be indicative of specific cell states (Guttman, 2009). A recent single cell RNA-sequencing study in fetal brain has confirmed these results in more mature neural tissue and shown that lncRNAs have higher expression in specific cell clusters than would it appear from bulk RNA-sequencing (Liu et al., 2016). Indeed, we find this to be the case within our own data where TrEx lncRNAs make up some of the top distinguishing genes between cell type clusters in week 2 cortical organoids (Fig. 4B-F & Table S4). In particular, we identified 8 TrEx lncRNAs that are expressed in single cell data and are predominantly expressed in one cell type. TrEx108 (CATG00000005887), TrEx2174 (RP11-314P15), TrEx2578 (NR2F2-AS1), and TrEx8168 (MIR219-2) are all most highly expressed in NE and absent in week 5 single cell data (Fig. 4C). TrEx5008 (RP11-71N10) and TrEx6514 (CATG00000085368) are predominantly restricted to the large RG cluster at week 2 (Fig. 4D). Interestingly, TrEx5008 appears to be transiently expressed in bulk RNA-seq data but is still highly expressed in a subset of RG in week 5 single cell RNA-seq data (Fig. 4B, 5D) indicating that some TrEx lncRNAs may be longer lived than what is implied by bulk RNA-sequencing and are in fact restricted to one cell subtype that may be overtaken by others as more cell types are generated. TrEx4039 (AC011306/M IR217HG locus) is lowly expressed in both NE and RG, but most concentrated in a portion of CR cells, indicating a role in differentiation, a specific cell subtype, or a cell state transition of CR cells (Fig. 4E). TrEx2819 (AC004158) had the most dispersed expression in both RG and NE cell clusters (Fig. 4F).

**Figure 4.**
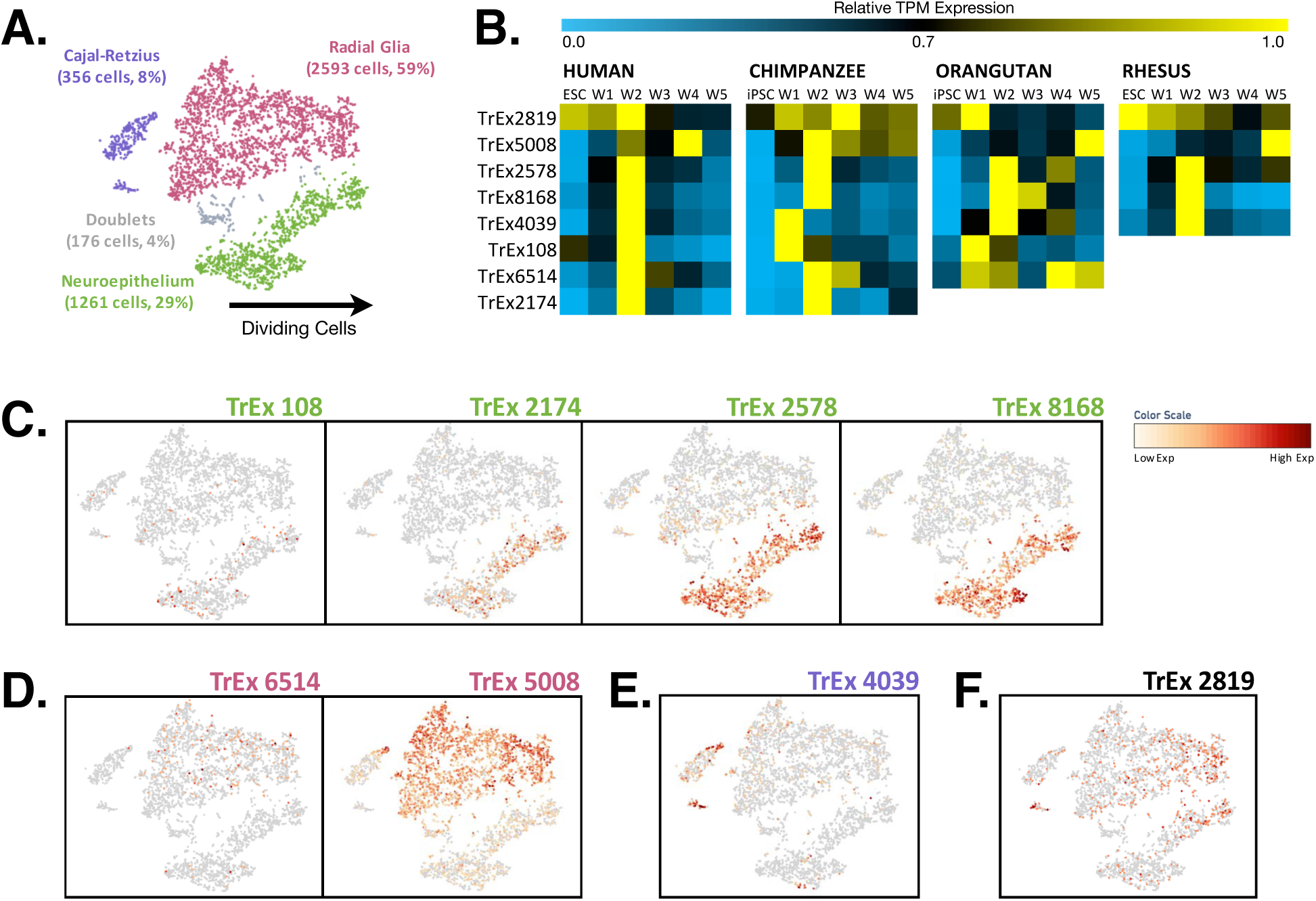
TrEx lncRNAs are implicated with specific cell subtypes by single cell RNA-seq. (A) A tSNE plot of manually curated cell types in week 2 single cell RNA-sequencing libraries. (B) A selection of 8 TrEx lncRNAs with transient expression patterns in human displayed in a heatmap showing their expression in species in which they are conserved by our metrics of gene structure and minimum expression. (C) TrEx108, TrEx2174, TrEx2578, and TrEx8168 were all expressed almost entirely within NE cells. (D) Both TrEx6514 and TrEx5008 were restricted mainly to RG. (E) TrEx4039 is expressed primarily in a subpopulation of the CR cell cluster. (F) TrEx2819 was expressed highly in both RG and NE, but appeared most highly expressed in dividing cells.

**Figure 5:**
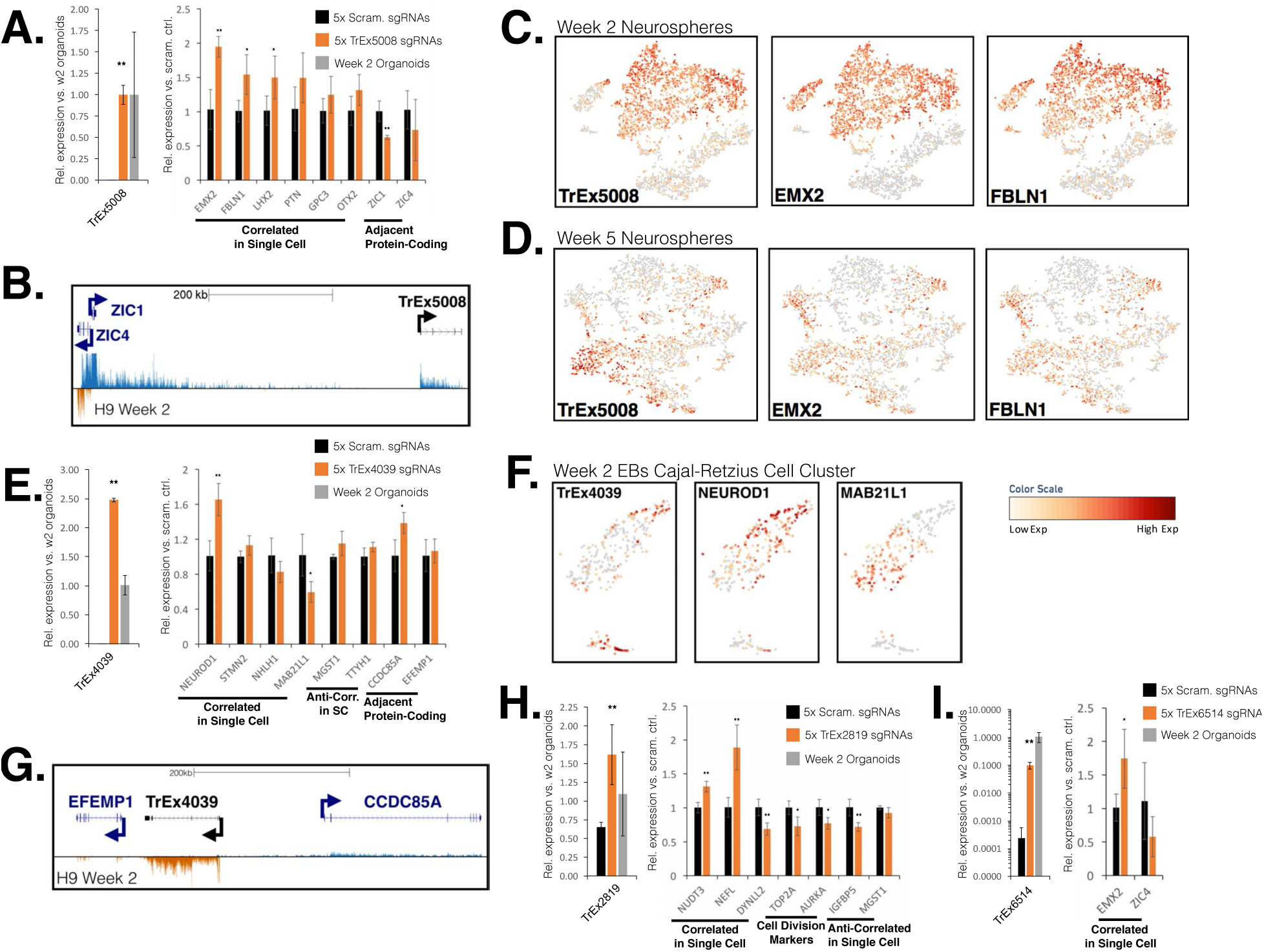
CRISPRa on TrEx lncRNAs up-regulates genes correlated in single cell RNA-seq. qRT-PCRs of target lncRNAs and their correlated genes from single cell RNA-sequencing upon CRISPRa of TrEx5008 (A), TrEx4039 (E), TrEx2819 (H), and TrEx6514 (I) in HEK293FT cells. * indicates p<0.05 and ** p<0.01 versus scrambled non-targeting controls. TrEx lncRNA expression in human week 2 cortical organoids is also shown to indicate normal expression levels of the gene. UCSC Genome Browser screenshots depicting a representative Cufflinks gene model of TrEx5008 (B) and TrEx4039 (G) and their nearest protein-coding genes. Coverage tracks from human week 2 bulk RNA-seq show positive strand reads represented in blue and negative strand in orange. t-SNE plots displaying expression of TrEx5008, EMX2, and FBLN1 from single cell RNA-seq at week 2 (C) and week 5 (D) of organoid differentiation show co-expression in human neurospheres. (F) t-SNE plots focusing on the CR cell cluster in week 2 organoid single cell RNA-seq display the expression of TrEx4039, NEUROD1, and MAB21L1.

### Out of context endogenous activation of lncRNAs regulates gene expression

Since many lncRNAs have been implicated in gene regulatory function in either *cis* (Leighton et al., 1995; Penny et al., 1996; Pandey et al., 2008; Zhao et al., 2008; De Santa et al., 2010; Kim et al., 2010; Orom et al., 2010; Wang et al., 2011) or *trans* (Rinn et al., 2007; Nagano et al., 2008; Pandey et al., 2008; Guttman et al., 2009; Khalil et al., 2009; Huarte et al., 2010; Kozoil and Rinn, 2010; Loewer et al., 2010; Zhao et al., 2010; Guttman et al., 2011; Hung et al., 2011) we assessed the potential gene regulatory function of TrEx lncRNAs by CRISPR activation (CRISPRa) using dCas9-VP64 to drive transcription from the endogenous locus in HEK293FT cells similar to Konermann et al., 2014. TrEx lncRNA candidates that were differentially expressed in bulk RNA-seq data, detectable by single cell RNA-seq, had little to no expression in HEK293FT cells, and appeared to be expressed from independent promoters in our bulk RNA-seq coverage were selected for endogenous locus activation. Priority was given to lncRNAs that were max expressed at week 2 by bulk RNA-seq or in the top 20 genes that distinguish best between NE, RG, and CR at week 2 (Fig. 4, S12). 4 or 5 CRISPR single-guide RNAs (sgRNAs) were designed 50 to 450bp upstream from each candidate TrEx lncRNA and co-transfected into HEK293FT with dCas9-VP64 to drive transcription from the endogenous locus. In all, we achieved activation of 8 TrEx lncRNAs ranging from 2.5-fold to 8600-fold activation over non-targeting scrambled sgRNA controls. Four of these were activated to a similar or higher expression level compared to bulk week 2 human cortical organoid RNA (Fig. 5, 6).

**Figure 6:**
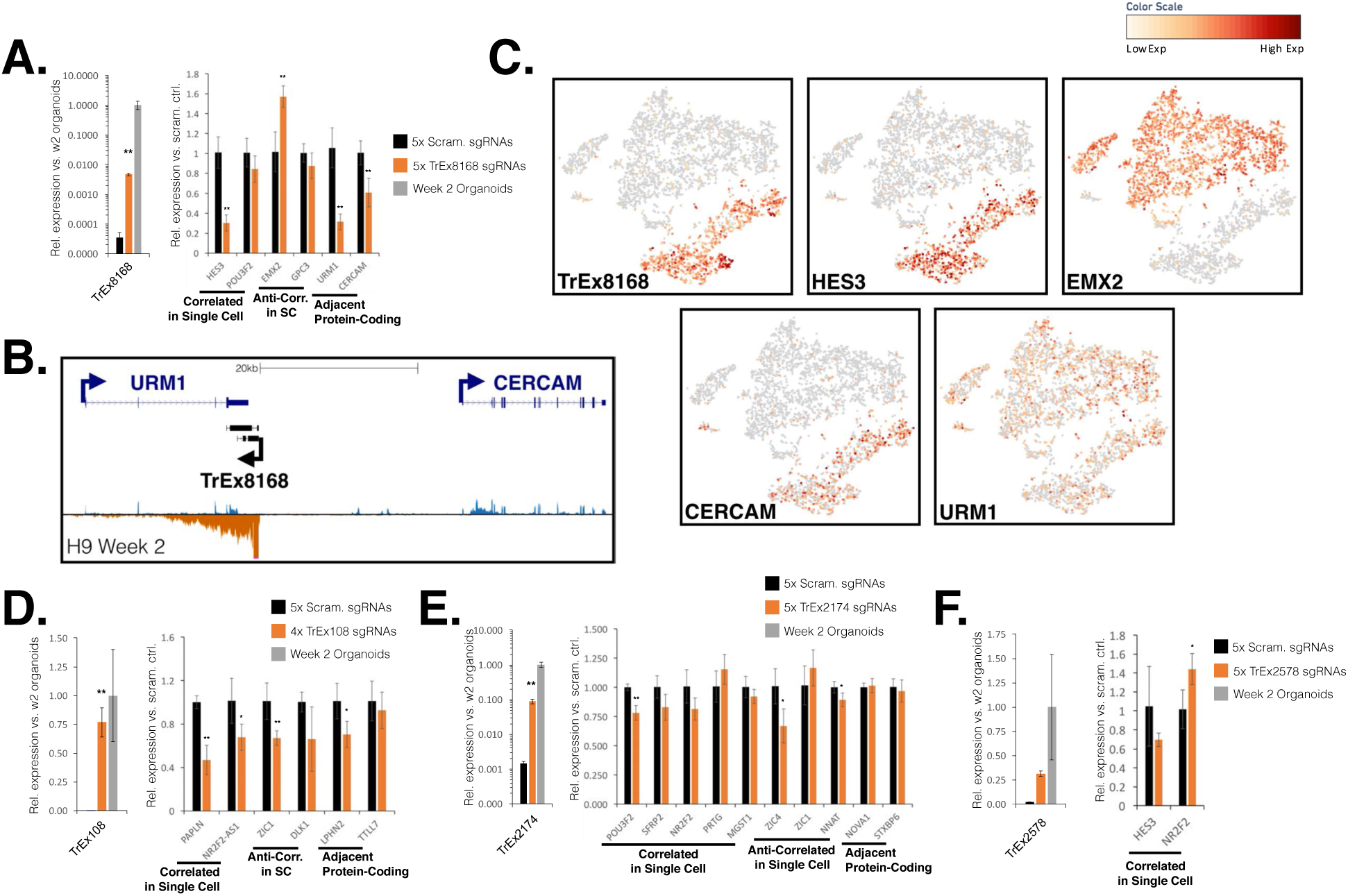
CRISPRa on NE associated TrEx lncRNAs. qRT-PCRs of target lncRNAs and their correlated genes from single cell RNA-sequencing upon CRISPRa of TrEx8168 (A), TrEx108 (D), TrEx2174 (E), and TrEx2578 (F) in HEK293FT cells. * indicates p<0.05 and ** p<0.01 versus scrambled non-targeting controls. TrEx lncRNA expression in human week 2 cortical organoids is also shown to indicate normal expression levels of the gene. (B) UCSC Genome Browser screenshot depicting a representative Cufflinks gene model of TrEx8168 and its nearest protein-coding genes, URM1 and CERCAM, is shown. Coverage tracks from human week 2 bulk RNA-seq were provided with positive strand reads represented in blue and negative strand in orange. (C) t-SNE plots displaying expression of TrEx8168, HES3, EMX2, CERCAM and URM1 from single cell RNA-seq of week 2 human organoids illustrate cell cluster association.

To assess the regulatory potential of these activated TrEx lncRNAs, we used quantitative reverse transcriptase PCR (qRT-PCR) to measure the expression of protein-coding genes that were correlated or anticorrelated with target TrEx lncRNA expression in single cell data as potential targets for lncRNA mediated regulation. A gene was considered correlated if either it was expressed in the same cell type in single cell RNA-seq or scored in the top 10 scores by Pearson correlation (Table S5, S6). Similarly, a gene was considered anticorrelated if it predominantly appeared in another cell type cluster or was in the bottom 10 scores by Pearson correlation (Table S5, S6). We tested expression changes of these genes upon CRISPRa of their respective TrEx lncRNAs compared to a set of scrambled sequence non-targeting guide controls (Fig. 5, 6). If a TrEx lncRNA locus was located within 200kb of a protein coding gene or its nearest neighbor exhibited cluster-specific expression in single cell RNA-seq, we also tested adjacent gene expression to look for *cis* regulatory action of the lncRNA or potential off-target effects of dCas9-VP64. We found that all 8 activated TrEx lncRNAs had reproducible effects on gene expression of at least one of their potential targets with a p-value of less than 0.05 in a two-tailed t-test. Though most showed only modest effects on distal gene regulation, 3 induced a clear 2-fold change or greater on gene expression in *trans*.

The two TrEx lncRNAs associated with RG cells, TrEx5008 and TrEx6514, generally activated genes that correlated well with their expression in single cell RNA-seq (Fig. 5A, I). TrEx5008 in particular exhibited significant activation of EMX2, increasing its expression 2-fold over non-targeting controls and a more modest, but consistent, effect on FBLN1 (1.5 fold) (Fig. 5A). Interestingly, TrEx5008 expression continues to track with EMX2 and FBLN1 in week 5 neurospheres within a subset of RG (Fig. 5C, D) further supporting an association with the same gene regulatory network. There is also a significant repression of the upstream gene ZIC1 (-1.6-fold) upon TrEx5008 activation (Fig. 5A, B). Since this gene is over 500kb upstream, this is unlikely to be due to direct dCas9-VP64 interference, and may instead be due to a local chromatin structure change, possibly influenced by a *cis* regulatory function of TrEx5008.

Though some of the activated TrEx lncRNAs had the effect of up-regulating positively correlated genes (i.e. TrEx5008, TrEx6514, TrEx4039, and TrEx2819) we found examples that had the opposite effect where genes positively correlated in single cell data were generally down-regulated upon activation (TrEx108, TrEx2174, TrEx8168). Curiously, these were all associated with NE and specifically expressed at week 2 as seen by both single cell and bulk RNA-seq. Specifically, activation of TrEx8168 significantly reduces *HES3* expression by about 70% and up-regulates the anti-correlated *EMX2* by almost 60% even with less than a tenth of the expression seen in week 2 organoids (Fig. 6A). TrEx8168 also seems to significantly disrupt the local transcriptional landscape, reducing the expression of its neighboring genes *CERCAM* by 40% and *URM1* by 68% even with less than 0.5% the level of expression as week 2 organoids (Fig. 6A & B). *URM1* appears to have a low dispersed expression pattern in single cell RNA-seq but *CERCAM* seems to be co-expressed in the same cell cluster as TrEx8168 (Fig. 6C). Despite this pattern, neither gene was found to be significantly correlated or anti-correlated by Pearson correlation with TrEx8168. TrEx108 also had drastic effects on the single cell expression associated gene, *PAPLN,* reducing its expression by 2.1-fold (Fig. 6D). Both of these TrEx lncRNAs may signal the exit from a neuroepithelial cell state as their expression disrupts gene regulatory networks involved in the maintenance of that state.

The CR associated lncRNA TrEx4039 was activated to a level 2.5-fold greater than bulk week 2 organoids and exhibited a 66% elevation in *NEUROD1* along with a decrease of about 40% in *MAB21L1* (Fig. 5E). While these effects are modest, they agree well with the heterogeneous distribution of *MAB21L1* and *NEUROD1* within the CR cell cluster (Fig. 5F). Indeed, expression of TrEx4039 is concurrent with *NEUROD1* but segregates from cells expressing *MAB21L1,* suggesting that TrEx4039 may contribute to the establishment or maintenance of a specific CR cell subtype or cell state.

TrEx2819 had the weakest change in expression versus the non-targeting control with about 2.48-fold activation, likely due to this transcript having low expression in unperturbed HEK293FT cells. However, this is a level of TrEx2819 expression similar to that observed in week 2 organoids (Fig. 5H). From the single cell data we hypothesized a role for TrEx2819 in cell division by its expression pattern in single cell RNA-seq, but we only observed a modest down-regulation of TOP2A and AURKA upon activation of TrEx2819. However, we did see a modest increase of two genes found by Pearson correlation, NUDT3 (1.3-fold) and NEFL (1.9-fold), and down-regulation of the correlated DYNLL2 (-1.5-fold). TrEx2819 action on cytoskeletal proteins may indicate a role in cell structure reorganization pre- or post-cell division.

We achieved relatively low activation of TrEx2174 (Fig. 6E) and TrEx2578 (Fig. 6F) compared to the expression in week 2 organoids, yet still saw effects on genes associated by single cell RNA-seq. TrEx2174 reduced the correlated POU3F2 by about 25% suggesting a role in the exit from NE cells. TrEx2578, despite being antisense to NR2F2, increased its expression by over 1.4-fold. However, these effects are more modest than those seen at the other loci, and may not suggest function.

## DISCUSSION

The RNA-Seq data generated in this study provides a valuable resource for comparative studies aimed at understanding human, chimpanzee, orangutan, and rhesus cortical development. These tissues provide insight into early differentiation stages that are largely inaccessible *in vivo* and could shed light on what makes great apes and humans unique from each other and from other species. Further, while human, chimpanzee, and rhesus cortical neuron differentiation has been studied with neural organoids, to our knowledge, we provide the first look at orangutan early cortical neuron differentiation events. This system allows us to look at equivalent time points during this important stage of development in a protocol that is robust in all of our pluripotent stem cell lines and species. Pairing weekly bulk RNA-seq across species with analysis of the cell type composition of these heterogeneous cultures by single cell RNA-seq in human provides additional insight into the context of expression events during the formation of these early neural cell types. Here we have gone deeper into a small subset of the functional gene regulatory elements influencing primate cortical development that might be found in these data. Many more could be discovered and studied by further analysis of this comprehensive RNA-seq data set.

The lncRNA field has been mired in controversy over the functional relevance of the tens of thousands of identified transcripts in human (Gingeras, 2012; Kowalczyk et al., 2012; Hon et al., 2017), with claims that, despite a few notable exceptions, most represent non-functional transcription from enhancer elements or spurious transcriptional noise (Ponjavic et al., 2007; Struhl, 2007; De Santa et al., 2010) due to their low sequence conservation across vertebrates (Wang et al., 2004; Babak et al., 2005; Pang et al., 2005; Ponjavic et al., 2007; Church et al., 2009; Kutter et al., 2012) or low levels of expression in bulk tissues (Cabili et al., 2011). It has also been suggested that tissue-specific lncRNAs are often less conserved than those expressed in multiple tissues (Ulitsky, 2016). We have shown here, however, that many human lncRNA transcripts expressed during the early stages of cortical neuron differentiation have structural conservation over great apes or to old world monkeys. Of the 2,975 lncRNAs expressed over our time course in human, 72% had conserved structure through chimp, 58% through orangutan, and 43% through rhesus. 51% were conserved in both great ape species and 31% had evidence of conserved structure in all species, much greater than the vanishingly small estimates of sequence conservation even of functional lncRNAs through mouse (Babak et al., 2005; Pang et al., 2006; Ponjavic et al., 2007). Striking among these transcripts were those that were specifically expressed at single time points in human cortical development, which we term TrEx lncRNAs. 386 of these TrEx lncRNAs were observed in human and had a remarkably conserved expression pattern in great ape species with at least 223 (58%) exhibiting a preserved TrEx pattern in chimpanzee or orangutan. While transient expression patterns seemed far less conserved than exonic structure, it is possible we are under-sampling relevant time points in each of our species for optimal detection of expression from these lncRNAs, especially considering most TrEx lncRNAs are primarily expressed at a single time point.

We found that many of these conserved transiently expressed transcripts were associated with specific cell types by single cell RNA-seq at the week 2 time point where there was a clear distinction between RG, NE, and CR cells (Fig. 4, Fig. S5). Using a strict definition of transcript conservation, requiring both gene intron boundary and expression pattern conservation between human and at least one other species, we set out to find transiently expressed lncRNAs associated with specific cell subtypes, reasoning that those have the highest likelihood of gene regulatory function. In all, 8 TrEx lncRNAs that appeared independently regulated in bulk RNA-seq with independent promoters from neighboring genes were selected for CRISPRa, allowing for detection of both cis and trans regulatory function. A recent study has shown that cellular context is vitally important for lncRNA function (Liu et al., 2017), so it is likely that the full functionality of these genes is significantly under-sampled by ectopic expression in our HEK293FT-based CRISPRa assay. Even still, we see effects on distal genes upon endogenous activation of these loci indicating robust gene regulatory function even out of its normal biological context, which warrants further study. All tested TrEx lncRNAs showed effects on distal gene regulation upon activation (Fig. 5 & 6), but only 3 showed effects of 2-fold change or greater on genes correlated or anticorrelated with their expression in single cell RNA-seq data (Fig. 5A & Fig. 6A,D).

We have demonstrated that lncRNAs expressed during early cortical neuron development are well conserved over short but significant evolutionary distances in exonic structure and, to a lesser extent, expression pattern. We found examples of transiently expressed lncRNAs with plausible roles in cell subtype specification and maintenance (TrEx5008), exiting progenitor states (TrEx108 and TrEx8168), and cell type establishment after differentiation (TrEx4039) using an assay that measures both *cis* and *trans* regulatory effects. If adapted to other tissues and timepoints, the approach outlined here provides a means to identify promising candidates for in-depth mechanistic studies to establish the functional roles for lncRNAs and deepen knowledge of gene regulation. The findings here only scratch the surface of our comparative bulk RNA and single cell RNA sequencing data sets. With thousands of expressed intergenic RNA elements, including hundreds of species-specific and cell-type specific elements, further interrogation of this time course data from human, chimpanzee, orangutan, and rhesus bulk RNA sequencing is likely to yield many more functional elements involved in primate cortical development and other processes, especially if effective filters are used. We find that by requiring a lncRNA to have a reproducible dynamic temporal expression pattern during development, tissue or cell type expression specificity, and evolutionary conservation of intron structure over short distances, e.g. between the great apes, we get a high rate of experimentally validated effects on transcription of other genes that are statistically associated with the given lncRNA in single cell expression data. By this approach, we may find that many more transiently expressed lncRNA loci are not just markers of cell type transitions, but serve their own functional roles in gene regulation.

## Acknowledgements

This work was supported by, CIRM Predoctoral Fellowship T3-00006 (ARF), CIRM Postdoctoral Fellowship TG2-01157 and Human Frontier Science Program Postdoctoral Fellowship LT000689/2010-L (FMJJ), CIRM Center of Excellence for Stem Cell Genomics (Stanford) GC1R-06673-A, CIRM Center for Big Data in Translational Genomics (SALK) GC1R-06673-B, and StemPath NIH/NIGMS R01 GM109031. David Haussler is an Investigator of the Howard Hughes Medical Institute. We thank Florence Wianny and Colette Dehay for generously providing LYON-ES1; Oliver Ryder for supplying Sumatran Orangutan fibroblast cell lines for reprogramming; Robert Diaz and Karen Shaff from Applied Stem Cell for their work preparing our chimpanzee iPSCs; Bryan King and Kristof Tigyi for assistance with animal handling for the teratoma assays; Susan Carpenter and Sergio Covarrubias for providing plasmids and expertise in designing CRISPRa experiments; Daniel Kim and Pablo Cordero for valuable discussions on experimental design and single cell RNA sequencing; Tom Nowakowski and Alex Pollen for advice in curating our single cell RNA-Seq clusters; Bari Nazario (UCSC Institute for the Biology of Stem Cells), Nader Pourmand (UCSC Genome Sequencing Center), and Ben Abrams (UCSC Life Science Microscopy Center) for their excellent technical support; Shana McDevitt at the UC Berkeley QB3 GSL for her expertise and support; members of the Haussler Lab for helpful discussions and support.

## Author Contributions

Conceptualization, A.R.F., S.R.S., and D.H.; Methodology, A.R.F. and F.M.J.J.; Investigation, A.R.F., F.M.J.J., A.P.R.P., A.M.R.-O., E.L., L.W., V.M., and J.L.R; Formal Analysis, A.R.F., I.T.F., M.H., and S.K.; Writing - Original Draft, A.R.F.; Writing - Review & Editing, S.R.S. and D.H.; Funding Acquisition, A.R.F., F.M.J.J., S.R.S., and D.H.; Supervision, S.R.S. and D.H.

## STAR METHODS

### iPSC generation

Primate primary fibroblasts were grown as adherent cultures in MEM Alpha (ThermoFisher) supplemented with 10% Gibco FBS (ThermoFisher) and 1% Pen-Strep (ThermoFisher). Integration-free chimpanzee induced pluripotent stem cells were produced at Applied StemCell from S008919 primary fibroblasts (Yerkes Primates, Coriell) by episomal reprogramming using the Y4 plasmid combination described in Okita et al., 2011. Integration-free Sumatran orangutan induced pluripotent stem cells were generated using the CytoTune 2.0 Sendai Reprogramming kit (ThermoFisher) from 11045-4593 primary fibroblasts obtained from the Frozen Zoo^®^ (http://institute.sandiegozoo.org/resources/frozen-zoo%C2%AE). Both chimpanzee and orangutan iPSCs were initially established on mouse embryonic fibroblasts with KSR-15 (KO DMEM/F-12 + 20% KOSR, 1% NEAA, 1% GlutaMAX, 1% Pen-Strep, and 0.1mM 2-mercaptoethanol supplemented with 15 ng/mL bFGF) media and were transferred to feeder-free conditions on Matrigel (Corning) with mTeSR-1 (Stem Cell Tech) for chimpanzee or vitronectin (ThermoFisher) with Essential-8 Flex (ThermoFisher) for orangutan. Pluripotency was confirmed by immunofluorescence staining of pluripotency markers, RT-PCR, teratoma assay, and karyotype (Fig. S1 & S2).

### Teratoma Assay

Mice were anesthetized by intraperitoneal injection with 100mg/kg ketamine (MWI Veterinary Supply). 2 subcutaneous injections of 1 to 5 million cells suspended in 30% Matrigel (Corning) were made in the dorsolateral or ventral lateral areas of NOD-SCID mice (NOD.CB17-Prkdc^scid^/NCrCrl, Charles River) similar to Prokhorova et al., 2009. Mice were observed for up to 12 weeks for the appearance of tumors in the injected areas. The animals were euthanized by cervical dislocation and teratomas were harvested, fixed in 4% paraformaldehyde, saturated in 30% sucrose in PBS, embedded in Tissue Freezing Medium™ (Triangle Biomedical Sciences), and frozen for cryostat sectioning. Sections of the tumors were stained with hematoxylin (Mayer’s Hematoxylin Solution, Sigma) & eosin (Eosin Y solution, Sigma) and analyzed for the generation of all three germ layers.

### Karyotyping

Chimpanzee and orangutan iPSC lines were confirmed to have a stable wild type 48/XX karyotype through at least passage 32 or 36, respectively (Fig. S1F & S2E). Karyotyping services were performed by Cell Line Genetics or the Coriell Institute for Medical Research.

### Cortical organoid generation

The Eiraku et al., 2008 protocol was optimized for use with human ESCs, rhesus ESCs, chimpanzee iPSCs, and orangutan iPSCs. Human H9 and rhesus LyonESC1 embryonic stem cells were cultured on mouse embryonic fibroblasts with KSR-8 media (KO DMEM/F-12 + 20% KOSR, 1% NEAA, 1% GlutaMAX, 1% Pen-strep, and 0.1mM 2-mercaptoethanol supplemented with 8ng/mL bFGF). Embryonic stem cells were manually lifted from MEF feeders and allowed to self-form into embryoid bodies on low attachment plates (Corning) in KSR media. Chimpanzee and orangutan induced pluripotent stem cells were grown in feeder-free conditions on matrigel (Corning) with mTeSR-1 (Stem Cell Tech) or vitronectin (ThermoFisher) with Essential-8 Flex media (ThermoFisher), respectively, and 10,000 cells per EB were aggregated using AggreWell-800 plates (Stem Cell Technologies) in Aggrewell media (Stem Cell Technologies) supplemented with 10 uM Y-27632 rock inhibitor (ATCC) and transferred to low attachment plates (Corning) on day 2. Both methods supplemented the respective media with 500ng/mL DKK1 (Peprotech), 500 ng/mL NOGGIN (R & D Systemes), 10 uM SB431542 (Sigma), and 1 uM Cyclopamine V. californicum (VWR) for the first 18 days of differentiation. The media was changed to Neuralbasal (Invitrogen) supplemented with N2 (Gibco) and 1 uM Cyclopamine on day 18. At this time, chimpanzee and orangutan neurospheres were also supplemented with 10ng/mL bFGF and 10ng/mL EGF to improve survivability in Neuralbasal media. After day 26, all cultures were grown in Neuralbasal/N2 media without any added factors. Total RNA was extracted at weekly time points for each species. This timeline was adjusted accordingly in rhesus, harvesting on days 6, 11, 17, 22, and 28, to account for differences in gestational timing (described below).

### Adjustment of rhesus macaque time points

When optimizing the ESC cortical neurosphere differentiation protocol for macaque ES cells, we noticed that macaque ESC colonies and neurospheres grow faster and form neural rosettes earlier than human. Indeed, macaque ESCs are reported to have a faster doubling time (~0.8 of hESCs; Amit et al., 2000; Fluckiger et al., 2006). Macaque also has a shorter gestation period and a predicted faster progression of neurodevelopmental events (Workman et al., 2013). Therefore, in order to most reliably compare the gene expression events that occur during human and macaque ESC cortical neurosphere differentiation we needed to adjust for the intrinsic difference in neurodevelopmental speed.

A time point adjustment was implemented based on semi-quantitative RT-PCR analysis comparing expression levels of *TBR1* and *CTIP2* over the course of ESC cortical neurosphere differentiation (Fig. S3). The best linear fit of human and macaque expression level dynamics is obtained when the relative timing for macaque cortical neurosphere differentiation is adjusted by a factor of 0.8, whereby human day 21 is most comparable to macaque day 17 and human day 35 is most comparable to macaque day 28 (Fig. S3A; right panel). Human, chimpanzee, and orangutan time points are indicated in weeks where the equivalent macaque time points are the time point most comparable to the corresponding human week: W1*, W2*, W3*, W4*, and W5* (days 6, 11, 17, 22, 28, respectively). No adjustment was found necessary for the other species in this study.

### Immunofluorescence Staining

Cortical organoids were fixed for 15 minutes in 4% paraformaldehyde and saturated in 30% sucrose prior to being embedded in Tissue Freezing Medium™ (Triangle Biomedical Sciences) and frozen for cryostat sectioning. Sections of 16-18uM were adhered to glass slides and fixed a second time in 4% paraformaldehyde for 10 minutes. Cells in 2-dimensional culture were grown on acid etched coverslips and fixed for 10 minutes with 4% paraformaldehyde prior to staining. Samples were incubated at 4°C in blocking solution (3% BSA and 0.1% Triton X-100 in PBS) for 4 hours. Primary antibody incubation was performed overnight at 4°C in blocking solution. Secondary antibody incubation was for 1-4 hours at room temperature in blocking solution. Samples were mounted with SlowFade^®^ Gold antifade reagent (Invitrogen). A full list of used primary and secondary antibodies can be found in the Supplemental Table S7.

### Primate Genome Alignment and Annotation

A progressive Cactus (Paten et al., 2011) whole genome alignment was generated between the human hg19 assembly, chimpanzee panTro4 assembly, orangutan ponAbe2 assembly, and rhesus macaque rheMac8 assembly. This alignment was used as input to the Comparative Annotation Toolkit (http://github.com/ComparativeGenomicsToolkit/Comparative-Annotation-Toolkit, Fiddes, et al, submitted) along with the FANTOM5 (Hon et al., 2017) lv3 annotation set.

This process projects the annotations from human to the other primates in the alignment. Subsequent filtering and post-processing produces a high quality comparative annotation set. RNA-seq obtained from SRA (http://www.ncbi.nlm.nih.gov/sra) were used to help guide the annotation process. The resulting annotations were used for the initial differential expression analysis below. Filtered FANTOM transcript models available on GEO (GSE106245): hg19.fantom.lv3.filtered.gtf.gz (human), panTro4.fantom.lv3.transmap.filtered.gtf.gz (chimpanzee), ponAbe2.fantom.lv3.transmap.filtered.gtf.gz (orangutan), rheMac8.fantom.lv3.transmap.filtered.gtf.gz (rhesus).

### RNA-Sequencing Analysis

Paired-end Illumina reads were trimmed from the 3’ end of read1 and read2 to 100x100bp for human and rhesus libraries and 80x80bp for chimpanzee and orangutan based on sequence quality. Bowtie2 v2.2.1 (Langmead et al., 2012) was used with the “--very-sensitive” parameter to filter reads against the repeatMasker library (Smit et al., 2015) for each respective species which were removed from further analysis. STAR v2.5.1b (Dobin et al., 2012) was used to map RNA-seq reads to hg19 (human, Genome Reference Consortium GRCh37, 2009), panTro4 (chimpanzee, CGSC Build 2.1.4, 2011), ponAbe2 (orangutan, WUSTL Pongo_albelii-2.0.2, 2007), and rheMac8 (rhesus macaque, Baylor College of Medicine HGSC Mmul_8.0.1, 2015) respective to the origin species. STAR was run with the default parameters with the following exceptions: -- outFilterMismatchNmax 999, --outFilterMismatchNoverLmax 0.04, --alignIntronMin 20, --alignIntronMax 1000000, and --alignMatesGapMax 1000000. STAR alignments were converted to genomic position coverage with the bedtools command genomeCoverageBed-split (Quinlan et al., 2010). Coverage for each gene in a gene model for its species was derived by summing the position coverage over all the exonic positions of the gene as defined by the annotation sets (GEO, GSE106245: hg19.fantom.lv3.filtered.gtf.gz [human], panTro4.fantom.lv3.transmap.filtered.gtf.gz [chimpanzee], ponAbe2.fantom.lv3.transmap.filtered.gtf.gz [orangutan], rheMac8.fantom.lv3.transmap.filtered.gtf.gz [rhesus]) for initial analysis. DESeq2 v1.14.1 (Love et al., 2014) was used to provide basemean expression values and differential expression analysis across the time course in each species. Total gene coverage for a gene was converted to read counts by dividing the coverage by N+N (100+100 for human and rhesus and 80+80 for chimpanzee and orangutan) since each paired-end NxN mapped read induces a total coverage of N+N across its genomic positions. Results are available at GEO, GSE106245: GSE106245_deseq_baseMean_allF2.txt.gz.

### LncRNA annotation analysis, structure conservation, and expression estimates

Cufflinks v2.0.2 suite (Trapnell et al., 2010; Trapnell et al., 2012) was used to assemble transcript predictions of potentially unannotated lncRNAs in each species and the Cuffmerge tool was used to combine these annotations with FANTOM5 transcripts. The resulting Cufflinks-assembled and merged transcript sets were then projected through the cactus alignment (Stanke et al., 2008) to each of the other three genomes. Guided by the Cufflinks annotation set in each genome, these projections from the other genomes were assigned a putative gene locus (GEO, GSE106245: hg19.cufflinks.filtered.gtf.gz [human], panTro4.cufflinks.filtered.gtf.gz [chimpanzee], ponAbe2.cufflinks.filtered.gtf.gz [orangutan], rheMac8.cufflinks.filtered.gtf.gz [rhesus]). Transcripts were assigned a lncRNA ID number based on overlap with block loci defined by co-expression of transcripts within 10kb at at least one time time point, expression from the same strand, uninterrupted by antisense transcription, and no overlap with known protein-coding genes (“lncRNA ID” column on tables in GEO, GSE106245).

RSEM v1.3.0 (Li and Dewey, 2011) was used to provide TPM expression values for these newly generated transcripts (GEO, GSE106245: GSE106245_hg19.RSEM.gene_expression.tsv.gz [human], GSE106245_panTro4.RSEM.gene_expression.tsv.gz [chimpanzee], GSE106245_ponAbe2.RSEM.gene_expression.tsv.gz [orangutan], GSE106245_rheMac8.RSEM.gene_expression.tsv.gz [rhesus]). Expressed lncRNAs were assessed using the homGeneMapping tool from the AUGUSTUS toolkit (Konig et al., 2016). homGeneMapping makes use of cactus alignments to project annotation features in all pairwise directions, providing an accounting of features found in other genomes. homGeneMapping was provided both the Cufflinks transcript assemblies as well as expression estimates derived from the combination of the week 0 to week 5 RNA-seq experiments in all four species. The results of this pipeline were combined with the above transcript projections to ascertain a set of lncRNA loci that appear to have human specific expression, human-chimp specific expression, great-ape specific expression, and expressed in all primates (Table S1). For this analysis, a locus was considered expressed in the current reference genome if one or more transcripts had RNA-seq support for every single one of its intron junctions, and considered expressed in another genome if the transcripts that mapped from that genome to the current reference had RNA-seq support for any of its intron junctions. All single-exon transcripts were filtered out to reduce noise. To eliminate the possibility of the specificity results being skewed by assembly gaps or alignment error, loci which appeared to have sub-tree specific expression were checked against the cactus alignment to ensure that there was a matching locus in each other genome. If a genome appeared to be missing sequence, then this locus was flagged as having incomplete information.

### 3’ Single Cell RNA-sequencing

Human H9 embryonic stem cells were grown on vitronectin with E8-Flex media (ThermoFisher). Neurospheres were aggregated and as described above above for chimpanzee and orangutan induced pluripotent stem cells. Single cell suspensions for 10X Genomics Chromium single cell RNA-sequencing were prepared with TrypLE (ThermoFisher) and handled according to the 10X protocol RevA (version 1 chemistry) for undifferentiated hESCs and week 5 cortical organoids and RevB (version 2 chemistry) for weeks 1 and 2 cortical organoids. Cell count, quality, and viability was assessed using Trypan Blue (ThermoFisher) on a TC20 automated cell counter (BioRad). Single cell suspensions were made aiming for 1500-3000 cells captured per library. The data was analyzed by CellRanger v1.2 (10X Genomics) using a custom annotation set based on FANTOM5 lv3 (Hon et al., 2017) (GEO, GSE106245: hg19.fantom.lv3.filtered.gtf.gz) and visualized using the Loupe Cell Browser v1.0.0 (10X Genomics). The gene by cell data matrices are available in supplemental data files in matrix market exchange format (http://math.nist.gov/MatrixMarket/formats.html) (GEO, GSE106245: sch9froz_wk01_af111_filtered_gene_bc_matrix.tgz [week 1 gene by cell matrix], sch9froz_wk01_af111_possorted.bam [week 1 position sorted BAM file] sch9froz_wk01_af112_filtered_gene_bc_matrix.tgz [week 1 gene by cell matrix] sch9froz_wk01_af112_possorted.bam [week 1 position sorted BAM file] ch9wild_wk00_af104_filtered_gene_bc_matrix.tgz [week 0 gene by cell matrix] ch9wild_wk00_af104_possorted.bam [week 0 position sorted BAM file] sch9wild_wk02_af106_filtered_gene_bc_matrix.tgz [week 2 gene by cell matrix] sch9wild_wk02_af106_possorted.bam [week 2 position sorted BAM file] sch9wild_wk02_af107_filtered_gene_bc_matrix.tgz [week 2 gene by cell matrix] sch9wild_wk02_af107_possorted.bam [week 2 position sorted BAM file] sch9wild_wk05_af102_filtered_gene_bc_matrix.tgz [week 5 gene by cell matrix] sch9wild_wk05_af102_possorted.bam [week 5 position sorted BAM file] sch9wild_wk05_af103_filtered_gene_bc_matrix.tgz [week 5 gene by cell matrix] sch9wild_wk05_af103_possorted.bam [week 5 position sorted BAM file]).

Cell clusters were identified and manually curated by the expression of canonical cell markers using a combination of graphical and K-means clustering from the CellRanger v1.2 pipeline (10X Genomics) as a guide (Table S4). Gene networks correlated with lncRNA expression were identified in two parallel methods in our week 2 organoid data sets. Within the Loupe cell Browser v1.0.0 (10X Genomics), cells expressing a target lncRNA were filtered and compared to all other cells by the “locally distinguishing genes” tool (Table S5). We also calculated the Pearson Correlation of every of the 8 TrEx lncRNAs against all other 59042 genes in the merged 10X expression matrix and selected the top 10 associated and anti-associated genes as potential gene regulatory targets (Table S6).

### CRISPRa assay

The CRISPR-activation (CRISPRa) assay was modified from Konermann et al., 2014. HEK293FT cells were cultured with DMEM+GlutaMAX (ThermoFisher) supplemented with 10% FBS without antibiotic. Each well of a 6-well plate was seeded with 500k cells and co-transfected at 60-70% confluence the next day using Xfect reagent (Takara) with dCas9-VP64_Blast (Feng Zhang, addgene #61425), MS2-p65-HSF1_Hygro (Feng Zhang, addgene #61426), and a combination of 5 custom guide RNAs per target in the custom plasmid 783 (gift from S. Carpenter, UCSC; guide sequences in Table S7) for a total of 7.5 g DNA in a ratio of 1:1:2, respectively. Transfected cells were selected at 24 hours by incubation with 2 g/mL puromycin until harvest. RNA was harvested at 48 hours after transfection using TRIzol reagent (ThermoFisher) and RNA was extracted using Direct-zol columns (ZYMO). Quantitect SYBR^®^ Green RT-PCR (Qiagen) was used with 50ng of total RNA per reaction, 4 replicates per condition (qPCR primer sequences provided in Table S7). Relative expression was calculated by ddCt normalized to HEK293FT transfection with non-targeting scrambled control guides.

